# Subharmonics and chaos in simple periodically-forced biomolecular models

**DOI:** 10.1101/145201

**Authors:** Evgeni V. Nikolaev, Sahand Jamal Rahi, Eduardo D. Sontag

## Abstract

This paper uncovers a remarkable behavior in two biochemical systems that commonly appear as components of signal transduction pathways in systems biology. These systems have globally attracting steady states when unforced, so they might have been considered “uninteresting” from a dynamical standpoint. However, when subject to a periodic excitation, strange attractors arise via a period-doubling cascade. Quantitative analyses of the corresponding discrete chaotic trajectories are conducted numerically by computing largest Lyapunov exponents, power spectra, and autocorrelation functions. To gain insight into the geometry of the strange attractors, the phase portraits of the corresponding iterated maps are interpreted as scatter plots for which marginal distributions are additionally evaluated. The lack of entrainment to external oscillations, in even the simplest biochemical networks, represents a level of additional complexity in molecular biology, which has previously been insufficiently recognized but is plausibly biologically important.

## 1 Introduction

Many unforced biochemical systems, such as pairs of mutually repressing genes, or phosphorylation/dephosphorylation cycles, can exhibit biologically important properties such as multistability and oscillations [1, 2, 3, 4, 5, 6, 7, 8]. The experimental observation of these behaviors helps one to distinguish among alternative models, and indicates the necessity of positive or negative feedback loops [9]. Biological observables exhibited in response to time-dependent forcing signals (i.e. “dynamic phenotypes”) can provide further insight into the structure of biological systems. Recent examples include scale invariance or “fold-change detection” [10, 11, 12], non-monotonic behavior under monotonic inputs [13], refractory period stabilization [14], and non-entrained solutions or “period skipping” when stimulii are periodic [14].

The paper [14] found experimentally, in a *C. elegans* odor sensing neuron, responses whose periods are roughly multiples of the period of an excitation signal (see sample time traces in Section SI-5), thus theoretically implying the presence of negative feedback loops, and went on to propose a circuit architecture that is capable of displaying the observed dynamic phenotype. These findings suggest the following theoretical question: what complicated dynamics can arise in the simplest biochemical systems, more generally, in response to a periodic input? Here we answer that question by showing that a negative feedback system motivated by [14], and also the “nonlinear integral feedback” circuit proposed in [10] for scale-invariance, can both exhibit a rich bifurcation structure and chaotic behavior in response to pulse-train excitations.

There is an extensive and deep literature that deals with the analysis of responses of nonlinear systems, and particularly oscillators, to periodic signals. Under the influence of external periodic environmental forcing, nonlinear systems can exhibit bifurcations leading to subharmonic responses and chaos. Such behaviors have been studied theoretically and experimentally, in squid axons [15], cellular circadian oscillations subjected to periodic forcing by a light-dark cycle [16], forced pendulums and other classical physical oscillators described by the van der Pol and the Duffing equations [17, 18, 19, 20, 21, 22, 23], and biochemical oscillators such as the “Brusselator” [24, 25]. Our contribution is to show that similar behaviors can be found already in two of the simplest nonlinear systems which appear in the current systems biology literature. Furthermore, and perhaps equally remarkable, our two systems are not rhythmic in the absence of periodic stimulation; quite the contrary, they have unique and globally asymptotically stable steady states when the input is constant. This complexity in ubiquitous systems that constitute components of typical signal sensing and transduction networks suggests a previously unrecognized hidden level of complexity in molecular biology, which is of plausible significance for biological function [14].

## 2 Setup

We first discuss the models to be considered, the type of periodic input, and the notion of chaos.

### 2.1 The two models

Our first example, motivated by the paper [14], where similar models appear, is a negative feedback system which consists of two species *X* and *Y* such that *X* enhances the production of *Y* and *Y* inhibits the production of *X.* The concentrations of *X* and *Y* at time *t* are denoted respectively by *x* = *x*(*t*) and *y* = *y*(*t*). The interaction terms are modeled by Michaelis-Menten kinetics, and both *X* and *Y* are subject to zeroth-order decay through a mechanism such as protease-mediated degradation. An external input *U*, with magnitude *u* = *u*(*t*) in appropriate units, triggers production of *X.* The equations are as follows:

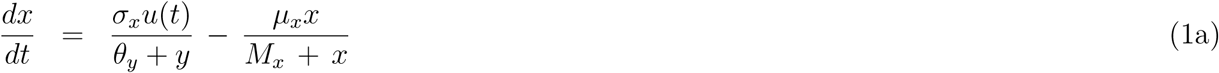

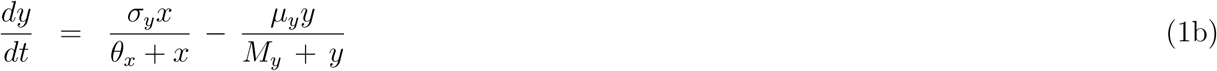

were we omit the argument *t* in *x* and in *y*, but leave it in the input in order to emphasize its time-dependence. All constants are assumed to be positive. A diagram of this model is in Fig. 1(a,c). with a general periodic input or a pulsed input respectively.

**Figure 1:**
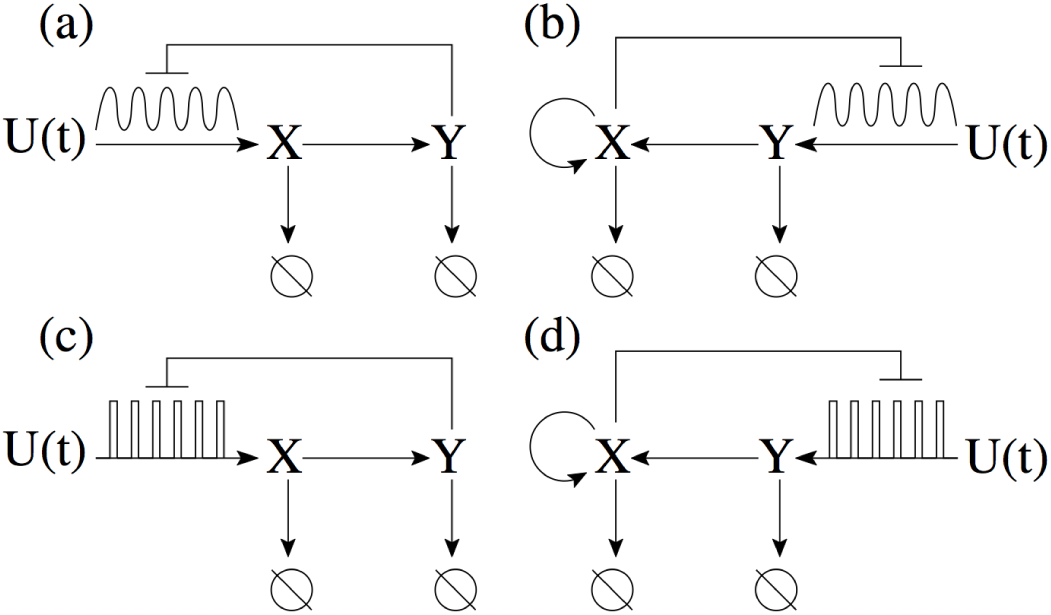
Simple biocircuits with an arbitrary periodic input and a negative feedback output. Left: negative feedback system. Right: FCD system. Top: arbitary periodic input. Bottom: pulse train input.

To emphasize that the bifucation behaviors are not an artifact of artificially chosen parameter values, we will also consider the special case of the model (1) in which all parameter values are unity (*σ*_*x*_ = *σ*_*y*_ = *θ*_*x*_ = *θ*_*y*_ = *µ*_*x*_ = *µ*_*y*_ = *M*_*x*_ = *M*_*y*_ = 1). We call this the “unity” model.

Our second example originates in the paper [10] (see [26] for more theoretical analysis). It is an integral feedback system consisting of a regulator species *X* and an output species *Y*, with equations (using again *x,y,u* for concentrations and inputs) as follows:

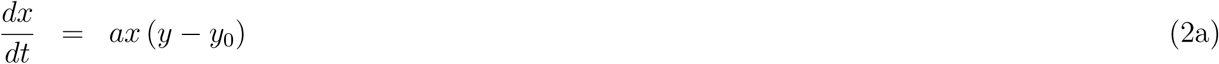

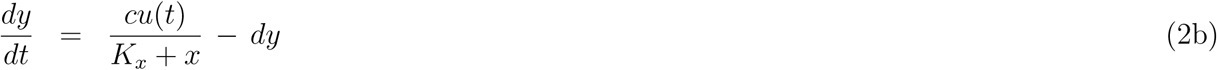

For a Michaelis-Menten constant *K_*x*_* ≪ 1, this system is perfectly adapting to constant inputs and has the fold-detection property, as discussed in [10]. A diagram of this model is in Fig. 1(b,d). with a general periodic input or a pulsed input respectively.

Both examples have the property that when *u*(*t*) is a constant there cannot be any periodic orbits and, furthermore, all solutions converge to a globally asymptotically stable steady state (with one minor exception for (1), explained in Section SI-1, that arises when some solutions become unbounded). For the system defined by (2), this fact was proved in [26]. For the system described by (1), this is discussed in Section SI-1.

### 2.2 Inputs

For simplicity, and to connect with the experiments in [14], we will consider *T*-periodic input functions *u*(*t*) (that is, *u*(*t*) ≡ *u*(*t* + *T*)) that consist of pulse trains of period *T*:

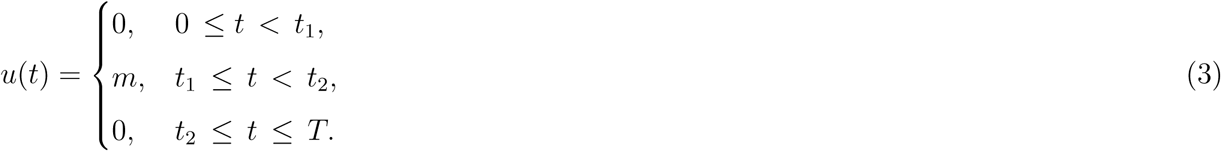

In (3), *t*_1_ = (*T* – ∆) /2, *t*_2_ = (*T* + ∆) /2, where *T* and ∆ are the period and support of the input function *u*(*t*), respectively. For the sake of (computational) simplicity, the support interval, [*t*_1_, *t*_2_], is centered at the midpoint of the interval [0,*T*]. We will analyze the effects of different choices of the amplitude parameter *m* or the period *T.*

### 2.3 Chaos

There are many definitions of chaos, the choice of which depends on the aspect of chaos to be emphasized [27, 28, 29, 30, 22, 31].

The definition of chaos appropriate for our work is based on periodic orbits and corresponds to the transition to chaos though period-doubling cascades [32, 33, 34]. Following [34], we say that the given dynamic system has a periodic-orbit chaos or a periodic orbit strange attractor, if it has infinitely many regular periodic saddles.

It is instructive to compare our biocircuit models with the Duffing’s equation, the second-order ordinary differential equation with a cubic nonlinearity describing a driven damped anharmonic oscillator, *x*″+ *kx*′+ *x*^3^ = *B* cos *ωt.* Here, the parameter *k* controls the damping, and the parameter *B* controls the amplitude of an external periodically varying driving force [17, 22]. Chaotic solutions of the Duffing’s equation were discovered as early as 1962–63 [20, 21, 23] and have been the subject of extensive research [35, 33, 18]. Detailed reviews of the Duffing equation’s fundamental properties can be found in [22, 31]. In the examples studied below, a strange attractor emerges via a cascade of period-doubling bifurcations, the transition to chaos also observed in the Duffing’s equation [33].

As already explained, when our biocircuits are subject to constant inputs, they admit a globally asymptotically stable state. However, when the external periodic force is allowed to act during a short period of time, very small relative to the period, chaotic behavior emerges (Fig. 2). This phenomenon is discussed in detail throughout the rest of the paper.

**Figure 2:**
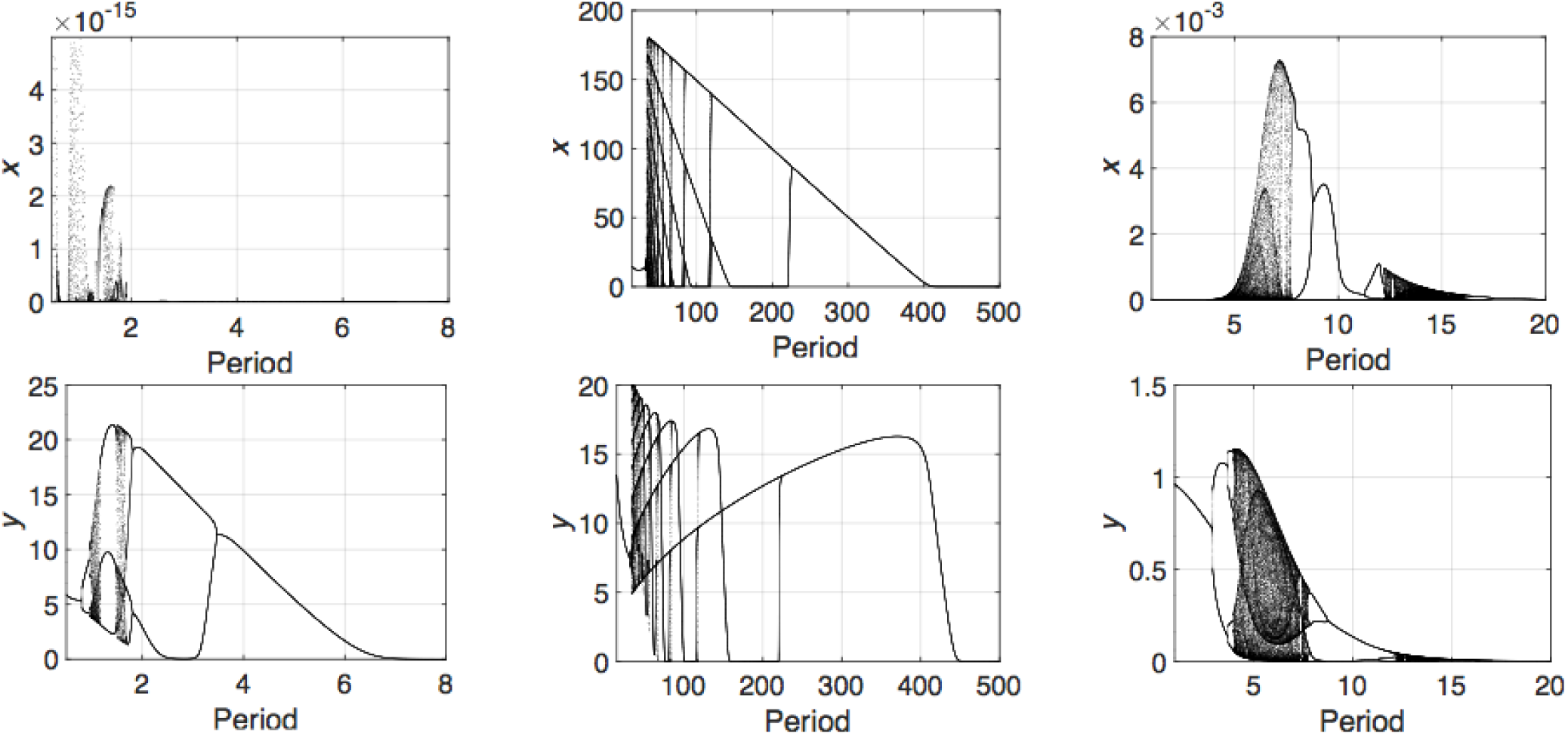
Bifurcation diagrams with respect to period T. The panels in the left column correspond to the model (1); the panels in the middle column correspond to the unity model obtained from the model (1) by setting all parameter values to one; and the panels in the right column correspond to the FCD model (2). The values of largest Lyapunov exponents for the attractors are shown in Fig. SI-2.1.

### 2.4 Shift Maps by Period *T*

The piecewise definition (3) of the input *u*(*t*) makes it inconvenient to numerically study bifurcations of limit cycles with respect to model parameters. It is more convenient to study the corresponding 2D-iterated maps, or shift maps by period *T*,

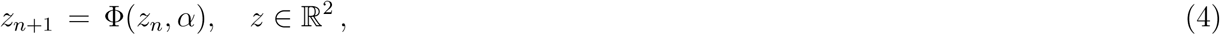

where Φ(*z_*n*_, α*) is the shift map by period *T* along the trajectories of the non-autonomous ODE given by (1) or by (2) respectively (recall that *T* is the (minimal) period of the external input *u*(*t*)). More precisely, we define Φ(*z, α*) = *φ*^*T*^ (*z, α*), where *φ*^*t*^(*z, α*) is the solution of the given ODE, that is, *φ*^*t*^ (*z, α*) = (*x*(*t, α*),*y*(*t, α*)), with the initial condition *z =* (*x,y*) at *t* = 0. The iteration (4) gives the value of *z*_*n+1*_ as a function of *z_*n*_* (*n* = 0, 1, 2, …), and *α* is the vector of model parameters. Note that a fixed point of the map (4) corresponds to the appropriate periodic solution of the ODE model considered.

Theoretical homotopy continuation [34] and numerical bifurcation approaches [36, 37], as implemented for instance in the powerful numerical bifurcation tool MatcontM [37], can then be applied to study numerically bifurcations of fixed and periodic points of the iterated map Φ(*z, α*) defined in (4). A technical requirement for these methods is that the map Φ(*z, α*) should be smooth with respect to state variable *z* and parameter *α.* This requirement is easy to verify, see Section SI-6.

## 3 Off-on-off chaos

Numerical analysis of the discrete trajectories of the maps (4) corresponding to the models described in Sect. 2.1 immediately reveals chaotic dynamics, in a wide range of model parameters.

Specifically Fig. 2 demonstrates examples of bifurcation diagrams generated from the corresponding iterated maps (4), when the values of the external period *T* are allowed to vary: (*a*) the leftmost column plots in Fig. 2 correspond to a strange attractor of the map (4) for the model (1) used with the fixed parameter values *σ*_*x*_ = *σ*_*y*_ = 10^4^, *θ*_*x*_ = 10, *θ*_*y*_ = 1, *µ*_*x*_ = 100, *µ*_*y*_ = 10, *M*_*x*_ = *M*_*y*_ = 1, *m* = 1.75, and ∆ = 10^−4^; (*b*) the middle column plots correspond to the map (4) for the unity model (1) (with all parameters equal to one) used with *m* = 2.0 x 10^4^ and ∆ = 10^−2^; and (*c*) the rightmost column plots correspond to a strange attractor of the map (4) for the FCD model (2) used with *a* = *Y*_0_ = *c*= *d* = 1.0, *m* = 10^−5^, and ∆ = 0.2.

We observe from Fig. 2 that for large values of the external period (that is, low frequencies, *ω* → 0), the dynamics in all the three models is asymptotically localized to a small vicinity of the unique globally stable steady-state, corresponding to oscillations with a very small amplitude. Indeed, large periods of external forcing allow the dynamics systems to relax to their globally stable steady states.

However, as soon as the period of the external input decreases, small amplitude oscillations develop into large-amplitude oscillations, followed by the transition to chaos via a period-doubling cascade. Remarkably, and counter-intuitively, while the period of the external input decreases, the period of periodic processes described by each periodically-forced model increases before each model becomes fully chaotic.

If the period *T* is further decreased, the chaotic dynamics is eliminated and is replaced again by regular periodic oscillations with the external period *T.* This backward transition, that is, “chaos → periodic oscillations,” can be explained by the Krylov-Bogoliubov-Mitropolsky (KBM) asymptotic theory [38, 39]. The theory predicts that asymptotically in the limit *ω* → ∞, the system dynamics becomes periodic with vanishing amplitude of oscillations. In this case, the high frequency small amplitude periodic dynamics can be approximated by a steady-state in the corresponding autonomous (averaged) system [40].

The strange attractors shown in Fig. 2 are called “off-on-off” attractors [34], because the chaotic dynamics disappears for small and large values of the bifurcation parameter, the external period *T* in this case, and exists only for intermediate values of the parameter. Examples of chaotic time-resolved solutions for the strange attractors (Fig. 2) with positive largest Lyapunov exponents (*λ*_max_ *>* 0) [41], are shown in Fig. SI-3.1.

To gain additional insight into the geometry of the strange attractors corresponding to the bifurcations trees (Fig. 2) and their complex time-resolved realizations (Fig. SI-3.1) we then plotted their respective phase portraits for the appropriate iterated maps (4) as discussed earlier. In the case of discrete trajectories generated by the map (4), it is convenient to interpret the phase portraits as scatter plots for which marginal distributions can be computed (SI-3). We observe from Fig. SI-3.1 and Fig. SI-3.3 that for the model (1) with the parameters shown, and also for the FCD-model (2), the discrete trajectories are localized along the axis *y*, respectively. However, in the special case of the unity model, the discrete trajectories stochastically jump between and stochastically move along several attractor loci (Fig. SI-3.1).

When instead of the period *T* the strength of the external input is taken as a bifurcation parameter, similar results are obtained. Bifurcations in the models leading to strange attractors in the case when the *m* is allowed to vary can be found in SI-4.

## 4 Power spectra and autocorrelation functions

In the theory of chaotic dynamic systems and discrete iterated maps, and specifically in [42] and references therein, the power spectrum is computed and used in order to distinguish periodic, quasiperiodic, and chaotic motions described by dynamical systems arising in a broad range of fields [30]. The power spectrum plotted for periodic or quasiperiodic trajectories has discrete peaks at the harmonic and sub-harmonics, whilst chaotic trajectories have a broadband component in their power spectrum. In order to illustrate that phenomenon for chaotic discrete trajectories of the iterated maps (4), we compute the power spectra and autocorrelation functions for the discrete trajectories corresponding to those values of parameters for which the plots in Fig. 2 (and Fig. SI-3.1) have been obtained.

To compute the power spectra we use Fast Fourier Transform (FFT) available from Matlab^©^ with *N* = 10^5^, the number of points used to compute the spectrum and autocorrelation functions (Fig. 3).

**Figure 3:**
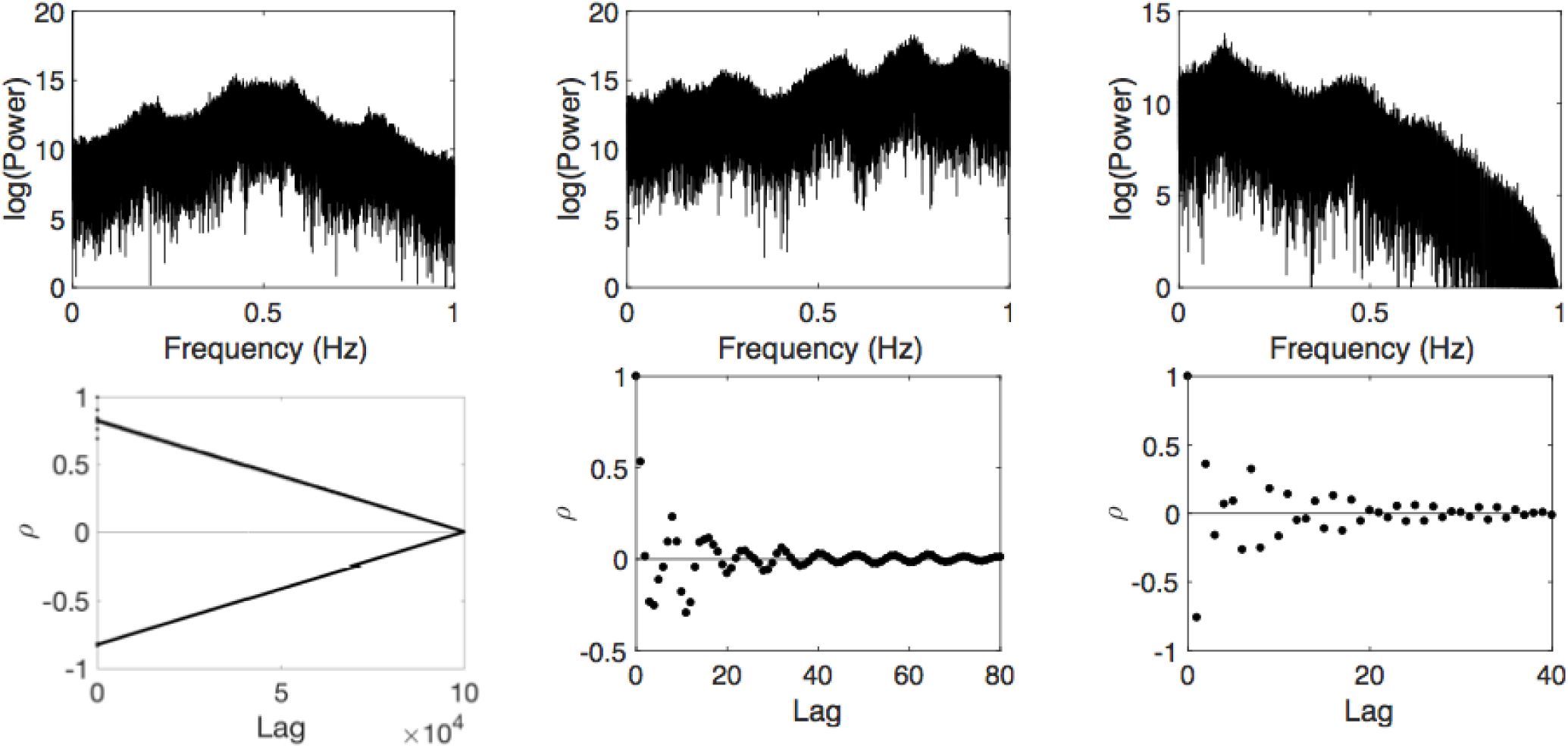
Power spectra and autocorrelation functions. Power spectra and autocorrelation functions are generated form the discrete chaotic trajectories of the corresponding iterated maps. The left column panels correspond to the map (4) for the model (1) with *T* = 1; the middle column panels correspond to the map (4) for the unity model with *T* = 40; and the right column panels correspond to the map (4) for the FCD model (2) with *T* = 5;

We can observe form Fig. 3 that the autocorrelation function for the model (1) converges to zero slowly, rapidly oscillating between positive and negative values, while the autocorrelation functions for the unity and FCD models vanish rapidly. This observation could be interpreted as saying that the model (1) has a larger capacity for memory than the other two models in terms of its remembrance of the past.

## 5 Myrberg-Feigenbaum Cascades

We continue our discussion of the bifurcation diagrams (Fig. 2) by analyzing the corresponding period-doubling cascades.

To being with, we note that Pekka Myrberg, a Finnish mathematician, was the first who discovered period-doubling cascades for periodic orbits with periods *p* x 2^*q*^,*q* = 1,2,3, …, for a variety of *p* values in a series of papers published in 1958–1963 [43, 44], reviewed in [34]. In the mid-1970s, Mitchell J. Feigenbaum discovered a remarkable universality for the period-doubling bifurcation cascades in 1D iterated maps [32]. Specifically, Feigenbaum discovered that if *d*_*k*_ is defined by *d*_*k*_ = *p*_*k*+1_ – *p*_*k*_, where *p*_*k*_ is the given map’s bifurcation parameter value corresponding to the *k-*th period doubling bifurcation with the period transition 2^*k*^ → 2^*k*+1^, then

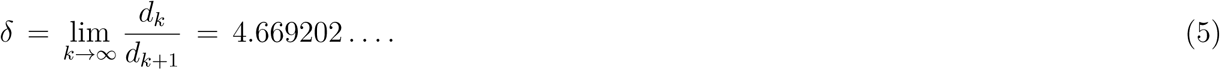

The number *δ*, known as the *Feigenbaum constant*, is as fundamental a quantity as the numbers *π* and *e*, in that it appears throughout the realms of science. The constant *δ* can be found not only in iterative maps but also in certain differential equations, for example, in the Duffing equation, as empirically shown in [33]. We next check if the corresponding bifurcation values of the parameters *T* and *m* also satisfy the universality law (5).

To carry out the corresponding computations systematically, we employed the command-line version of MatcontM [37]. Specifically, we first used MatcontM to compute the Feigenbaum (bifurcation) tree (Fig. 4), leading to the rapid accumulation of regular saddle periodic points which eventually form an infinite countable set around the parameter value *T*\* (the left panel) or *m*\* (the left panel), correspondingly. Here, *T*\* (or *m\**) corresponds to the Feigenbaum constant in the limit, *T*_*k*_ → *T*\* (or *m*_*k*_ → *m*\*) and *δ*_*k*_ → *δ*, as *k* → ∞ (Table 1).

**Figure 4:**
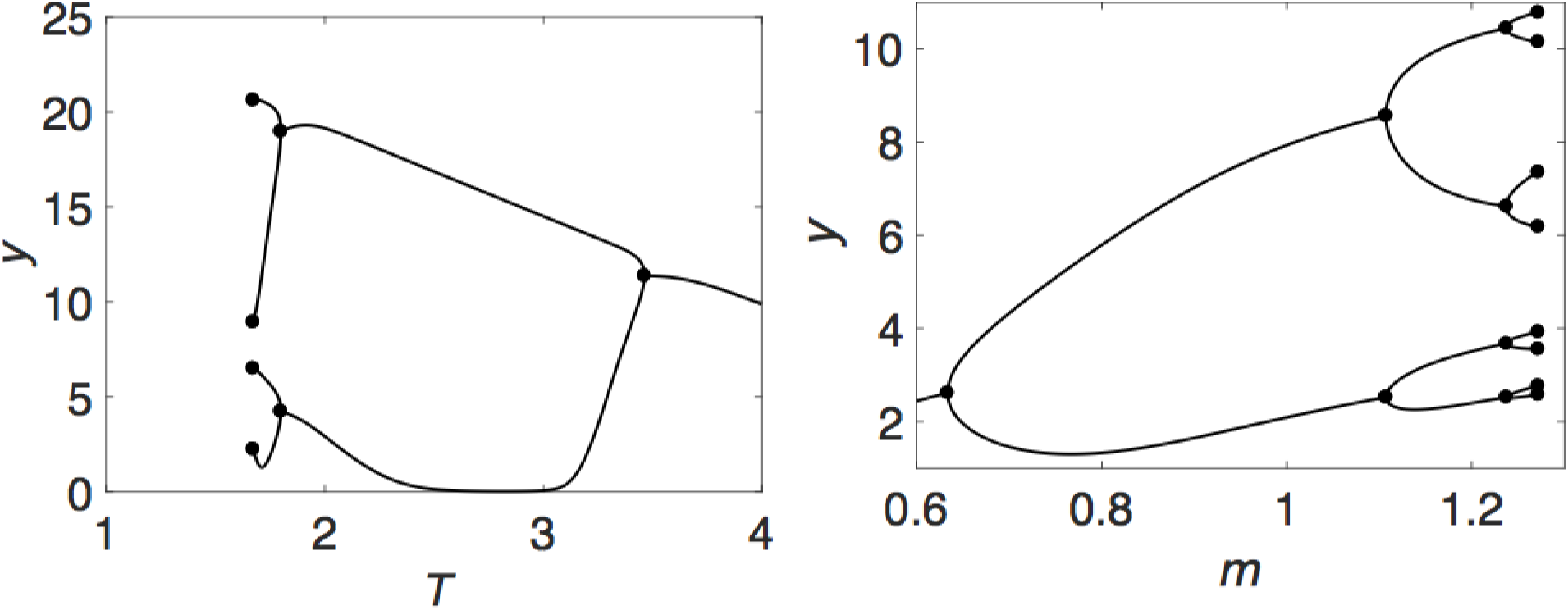
A Feigenbaum period-doubling tree. Because of a very fast accumulation of the bifurcation values of the corresponding parameters, *T*_*k*_ and *m*_*k*_, around their respective limits (Table 1), only very well visible onsets of the bifurcation trees are shown: The left panel corresponds to the bottom leftmost panel in Fig. 2, while the right panel corresponds to the bottom leftmost panel in Fig. SI-4.1.

**Table 1:** Numerical approximation of the Feigenbaum constant *δ.*

| $k$ | $T_k$ | $\delta_k$ | $m_k$ | $\delta_k$ |
| --- | --- | --- | --- | --- |
| 1 | 3.46269 | — | 0.63353 | — |
| 2 | 1.79848 | — | 1.10720 | — |
| 3 | 1.67155 | 13.11112 | 1.23735 | 3.63917 |
| 4 | 1.66440 | 17.74194 | 1.27174 | 3.78548 |
| 5 | 1.66345 | 7.52016 | 1.27984 | 4.24533 |
| 6 | 1.66326 | 5.13666 | 1.28161 | 4.55884 |
| 7 | 1.66322 | 4.72916 | 1.28200 | 4.64452 |

We find from Table 1 that while the sequence {*T*_*k*_} converges slowly, the sequence {*m*_*k*_} converges rapidly to its limit and its rate of convergence approximates the Feigenbaum constant very well.

To clarify our computational approach, we have to note that the default setup of MatcontM assumes (implicitly) that the map Φ(*z, α*) is given by explicit analytic expressions (formulas), since it uses algorithmic differentiation to compute Poincare normal form coefficients [45]. Because our map Φ (*z, α*) is defined implicitly through a number of numerical integration steps, we disabled the algorithmic (also called automatic or symbolic) differentiation (`adtayl`) feature by setting ‘`AutDerivative`’ = 0.

To ensure robustness of all numerical commutations, we used the MATLAB^©^ ode45, ode15s, and ode23s solvers with tight values of ‘`RelTol`’ = 10^−8^ and ‘`AbsTol`’ = 10^−10^, and, additionally, with ‘`RelTol`’ = 10^−10^ and ‘`AbsTol`’ = 10^−12^. To obtain all bifurcation values of both *T*_*k*_ and *m*_*k*_ with at least five significant digits, we computed the corresponding values two times by setting (*i*) ‘`FunTolerance`’ = 10^−6^ and 10^−8^, (*ii*) ‘`VarTolerance`’ = 10^−6^ and 10^−8^, and (*iii*) ‘`TestTolerance`’ = 10^−5^ and 10^−7^ inside MatcontM.

## 6 Multiple cascades

We next proceed with the discussion of the bifurcation diagrams shown in the middle column of Fig. 2 computed for the unity model. Specifically, intervals in the values of the parameter *T* with chaotic and regular dynamics interchange (Fig. 5, the left column panels). A similar phenomenon holds for the bifurcations with respect to the parameter *m* characterizing the input strength (Fig. 5, the right column panels and Fig SI-4.1). Fig. 5 shows the onset of chaos presented in the middle column panels of Fig. 4 and Fig SI-4.1, respectively. We next provide some more technical details, in order to obtain better insight into our numerical observations. We do this only for the parameter *T.*

**Figure 5:**
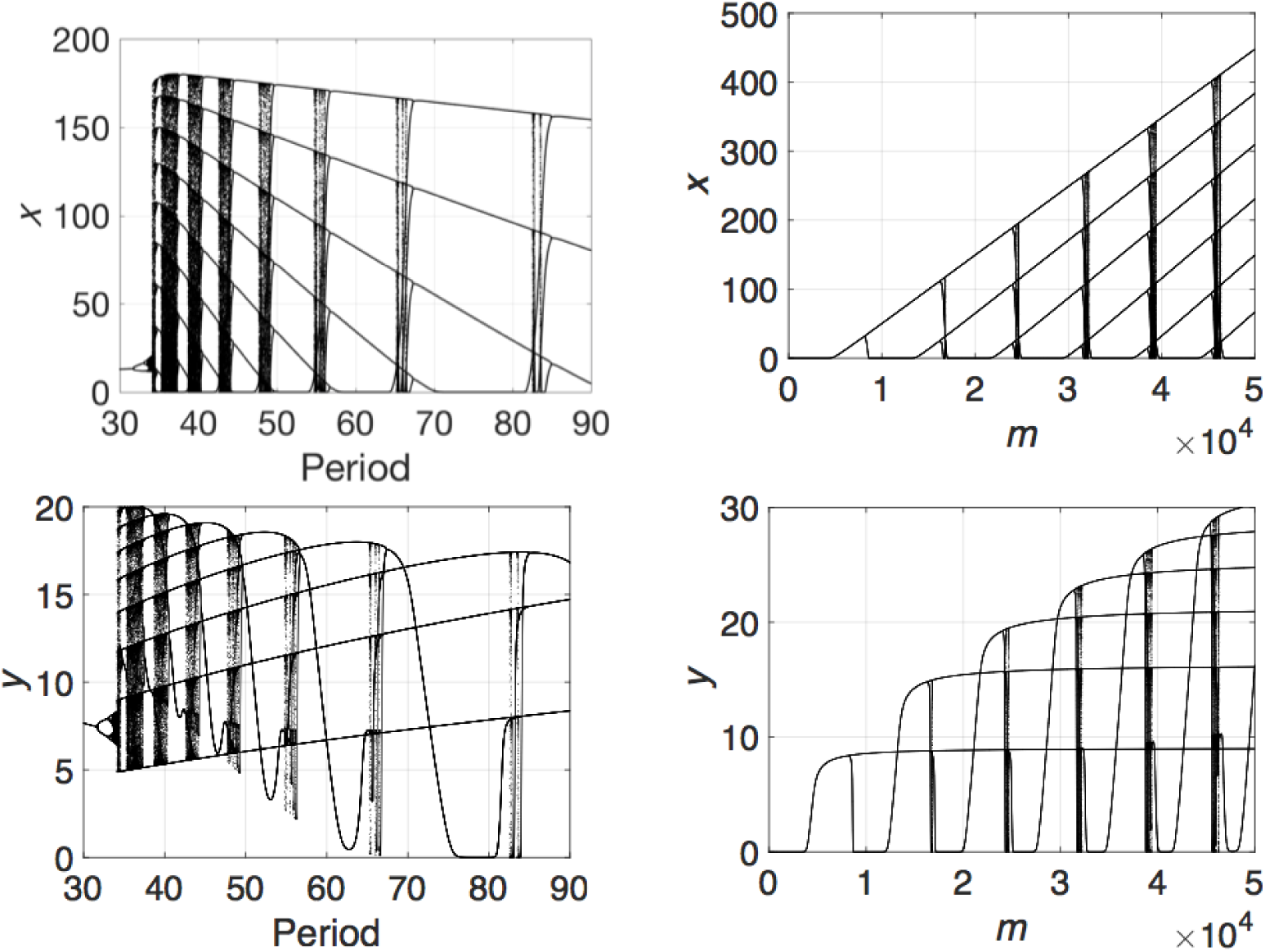
Multiple cascades in the unity model.

First, for each discrete value of the parameter *T*_*k*_, where *T*_*k*_ = *T*_0_ + *k*∆*T, T*_0_ = ∆*T* = 1, and *k* = 0,1,2, …, we generated the solution of the unity model over large time intervals with the initial conditions *x*(*0*) = *y*(*0*) = 10^−6^.

Second, using the solution obtained as a time series, we computed the *λ*_max_ and checked if it is equal to or is greater than zero.

Third, we found that chaotic intervals (*λ*_max_ > 0) in *T* are interchanged with intervals corresponding to regular periodic solutions (*λ*_max_ = 0).

The behavior of the unity model turned out to be so complex that it was impossible to employ MatcontM to carry out a detailed numerical bifurcation analysis similarly to that completed for the model (1), and discussed in Sect. 5. Despite the complexity of the observed dynamics (Fig. 5), we emphasize here that this model is exactly the type of very complex dynamical system for which the theory developed in [34] can applied to attain insight into the origin of the model’s chaotic behavior. Due to the theory [34] and our numerical evidence, we have a strong belief that multiple period-doubling cascades contribute to the formation of the strange attractor in both cases: when the values of *T* are allowed either to increase or to decrease, starting with those values of *T* for which a numerically verified globally stable unique periodic orbit exists. The case when the values of the parameter *T* are allowed to increase is shown in Fig. 5. The multiple cascades (Fig. 5) also lead to the formation of an infinite countable set of saddle periodic points which give rise the emergence of chaos in the unity model in accordance with the theory in [34].

## 7 Unbounded Chaos

We complete our discussion of chaos emerging in these simple periodically forced models with the case in which the FCD model has a strange attractor which appears to exist for unbounded values of the bifurcation parameters *T* and *m* as verified in our intensive numerical computations (Fig. 6 and Fig. SI-4.1).

**Figure 6:**
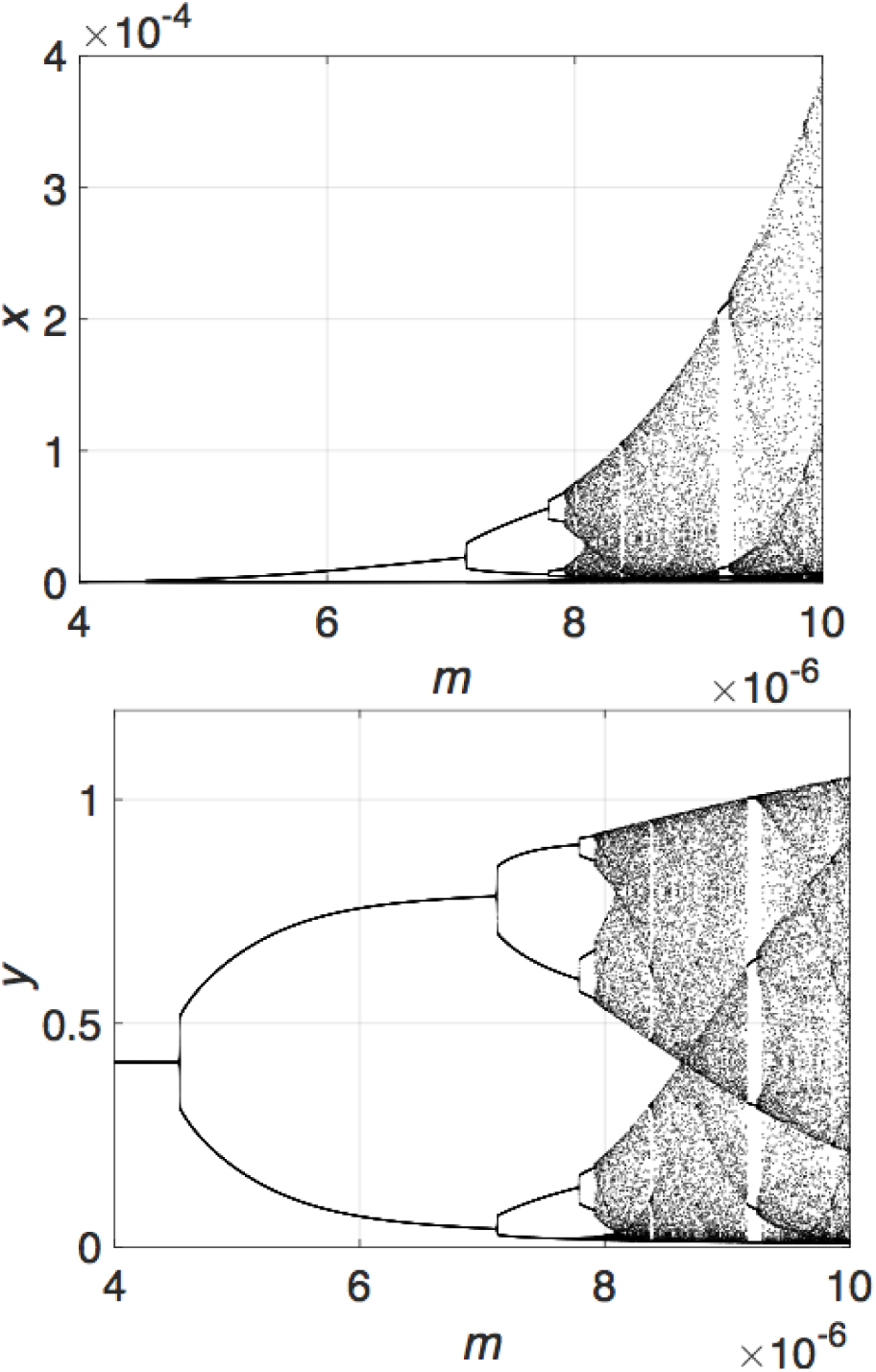
Unbounded chaos cascade in the FCD model with *T* = 5.

The plots (Fig. 6) show the onset of the chaos presented in Fig. SI-4.1 (the right column panels), where the latter is computed for a larger interval of the values of the parameter *m*. Both figures are computed for the same values of the fixed parameters.

We observe form Fig. 6 and Fig. SI-4.1 (the right column panels) that while the magnitude of chaotic changes in the *y*-variable stays bounded, the magnitude of chaotic changes in the *x*-variable grows monotonically as the values of the bifurcation parameter *m* increase (Fig. 6, the top panel), see also Fig. SI-4.1 (the right column panels). The complex dynamics within such strange attractors is called unbounded chaos [34]. Moreover, the theory predicts the existence of a cascade leading to the formation of strange attractors with unbounded chaos [34].

## 8 Discussion and conclusion

We showed that simple biochemical systems commonly seen in models of sensing and signal transduction pathways are able to exhibit a rich bifurcation structure into subharmonic oscillations, and even chaotic behavior, in response to periodic excitations.

The appearance of subharmonic responses is by no means an automatic property of cellular biochemical systems, however. For example, models of processes involved in gene transcription [46] and mRNA translation [47] can be shown to display the opposite behavior, namely entrainment, which means that all solutions have the same frequency as the forcing periodic signal. More generally, the synchronization of oscillators to external signals whose magnitude is large enough to enter the “Arnold tongue” insures that solutions will have the same frequency as the input [48]. As pointed out in the recent paper [49], entrained responses of biological systems play a key regulatory role in organisms [50, 51, 52].

While the behavior that we uncovered theoretically is consistent with the non-entrained responses seen experimentally in [14], it is virtually impossible to mathematically prove that an experimentally observed behavior is chaotic, or even perfectly subharmonic, especially in molecular biology, where noisy and relatively low precision measurements are the rule. Nonetheless, this work can serve as an indication that a lack of entrainment, complicated bifurcation structure, and chaotic behavior even in some of the simplest biochemical models, need not necessarily appeal to randomness or complex hidden regulatory pathways.

## Acknowledgements

Dr. Johannes Larsch generously shared the experimental data in Figure SI-5 with us.

## Supporting Information

### SI-1 Global dynamics of the model (1) with constant imputs

Consider the system described by (1), with a constant input *u*(*t*) = *u*_0_:

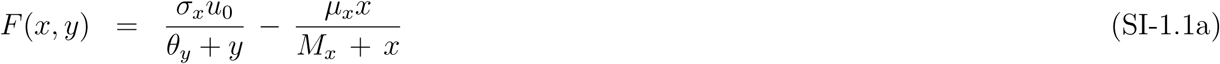

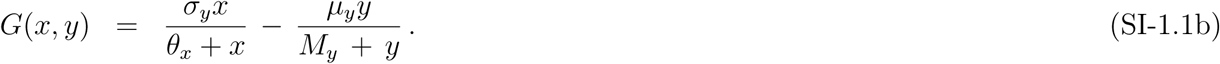

We first note that a steady state, if it exists, is unique (clear because of the increasing or decreasing character of the functions of *x* and *y*). An analysis of nullclines reveals that there is only one case in which solutions may become unbounded, see Figure SI-1.1, and our examples never treat that case.

**Figure SI-1.1:**
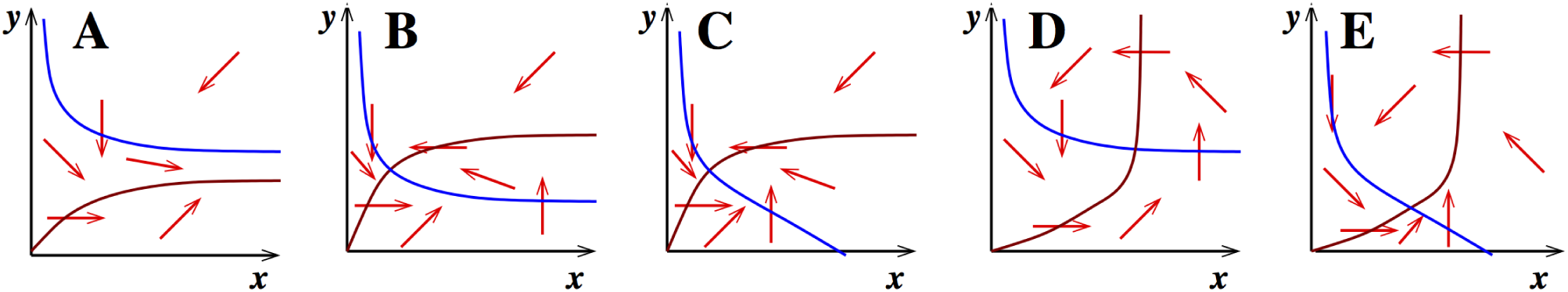
Possible phase planes. The blue curve represents the *x*-nullcline, that is, the locus of 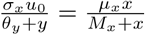, and the brown curve represents the *y*-nullcline, that is, the locus of 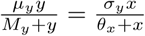. Arrows indicate directions of movement. In cases B-E, solutions remain bounded. In case A, solutions may diverge. Case A occurs when both of the following conditions hold: (i) *µ*_*y*_ > *σ*_*y*_ and (ii) *σ*_*y*_*M*_*y*_/(*µ*_*y*_ – *σ*_*y*_) < (*σ*_*x*_*u*_0_/*µ*_*x*_) – *θ*_*y*_. (These conditions characterize that case when both nullclines have have a positive limit as *x* → ∞, and they do not cross.) Note that we picked parameters so that (i) fails: *µ*_*y*_ = 10, *σ*_*y*_ = 10^4^ (or both equal to 1 in the “unity” model).

Once that boundedness if solutions is established, the Poincaré-Bendixson Theorem [53] insures that every solution converges to the unique equilibrium, unless there are periodic solutions or heteroclinic (including homoclinic) connections. However, periodic solutions and connections are ruled out by the Bendixson criterion [53], as follows. Consider the vector field *V*(*x, y*) = (*F*(*x, y*), *G*(*x, y*)), corresponding to the model in equations (SI-1.1). The Bendixson’s criterion states that if div*V*(*x, y*) ≠ 0 for all (*x,y*) ∈ *D*, then then the vector field *V*(*x,y*) does not have a closed orbit or heteroclinic connections in *D*, where *D* is any simply connected region of ℝ^2^, *D* ⊆ ℝ^2^, and

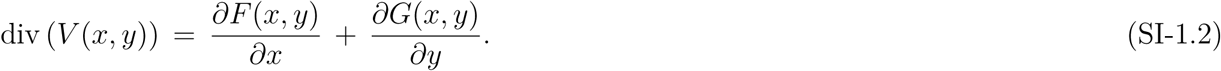

To show that div*V*(*x,y*) ≠ 0, ∀ (*x,y*) ∈ *D* = ℝ^2^, we simply compute:

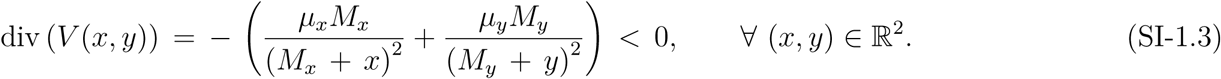

### SI-2 Numerical evaluation of largest Lyapunov exponents

**Figure SI-2.1:**
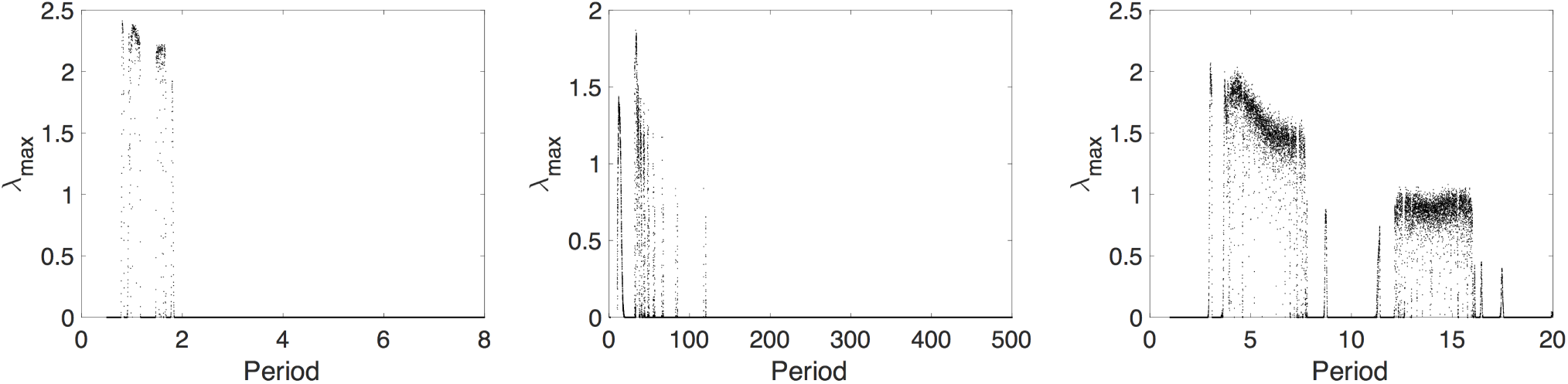
Largest Lyapunov exponents *λ*_max_ depending on period *T*. The panels corresponds to the panels from Fig. 2 in the main text. Positive values *λ*_max_ *>* 0 characterize chaotic solutions, while zero values *λ*_max_ = 0 are associated with periodic solutions of the corresponding models under parameter values given in Sect. 3 of the main text.

### SI-3 Examples of chaotic solutions

**Figure SI-3.1:**
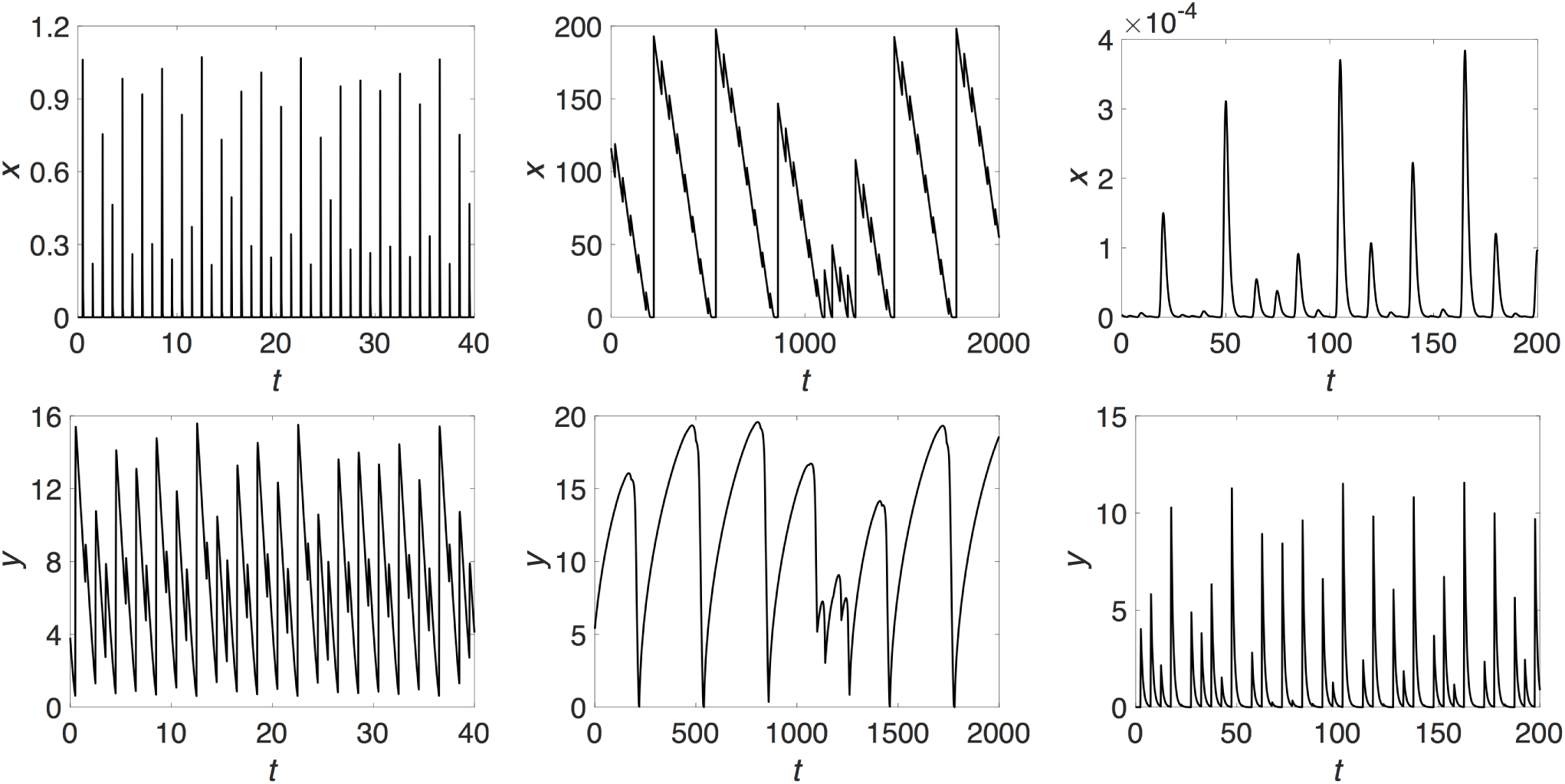
Chaotic solutions. The panels in the left column correspond to the model (1) with *T* = 1.0 (which results in *λ*_max_ ≈ 2.02); the panels in the middle column correspond to the unity model with *T* = 40 (which results in *λ*_max_ ≈ 0.86); and the panels in the right column correspond to the FCD model (2) with *T* = 5 (which results in *λ*_max_ ≈ 1.03). All other fixed parameter values are given in Sect. 3. (In the top left panel, *x*(*t*) decays extremely fast, hence the “spike” look of the plot.)

### SI-3 Scatter plots and marginal distributions

**Figure SI-3.1:**
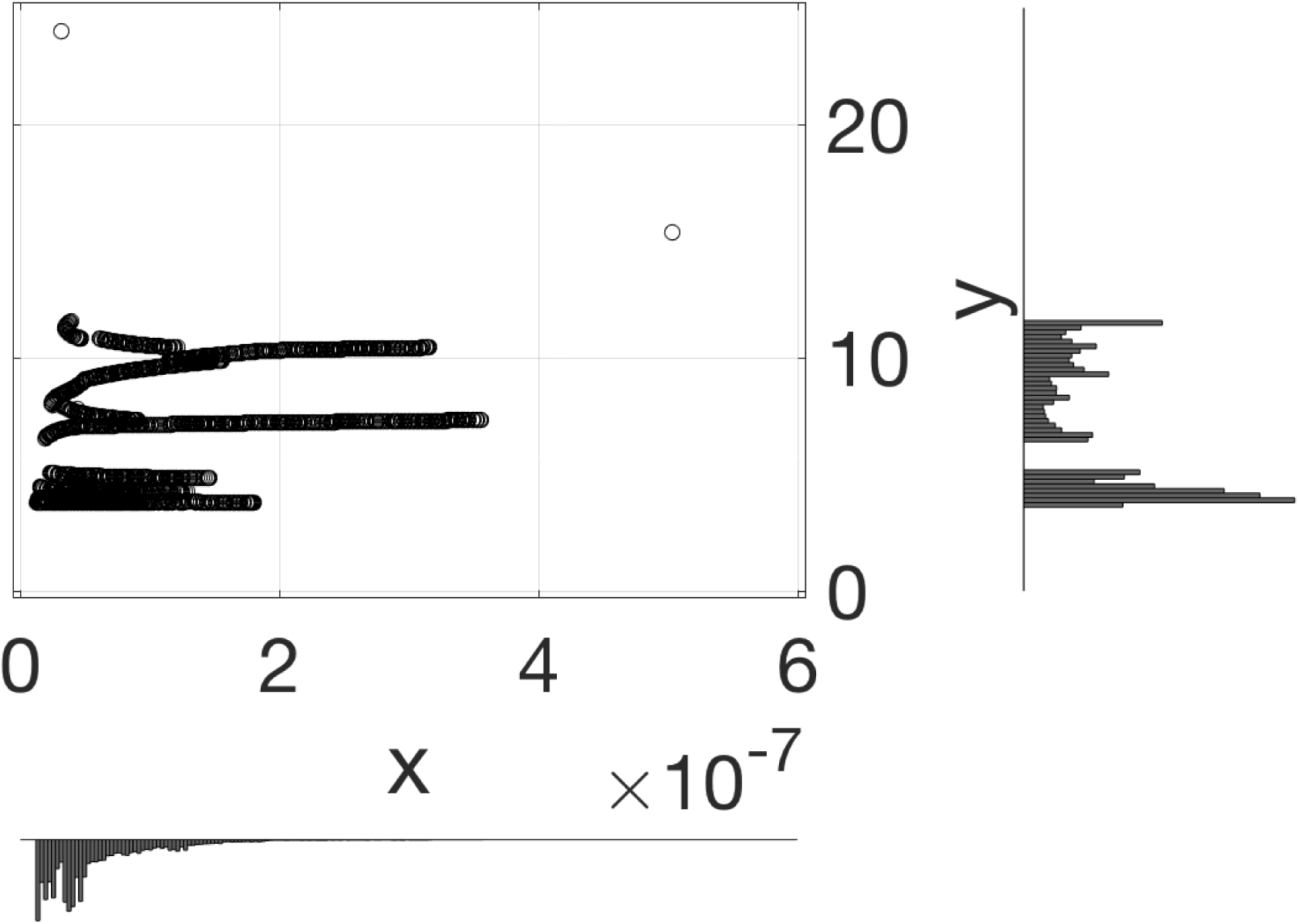
Scatter plot and marginal distributions for the model (1) with *T* = 1.

**Figure SI-3.2:**
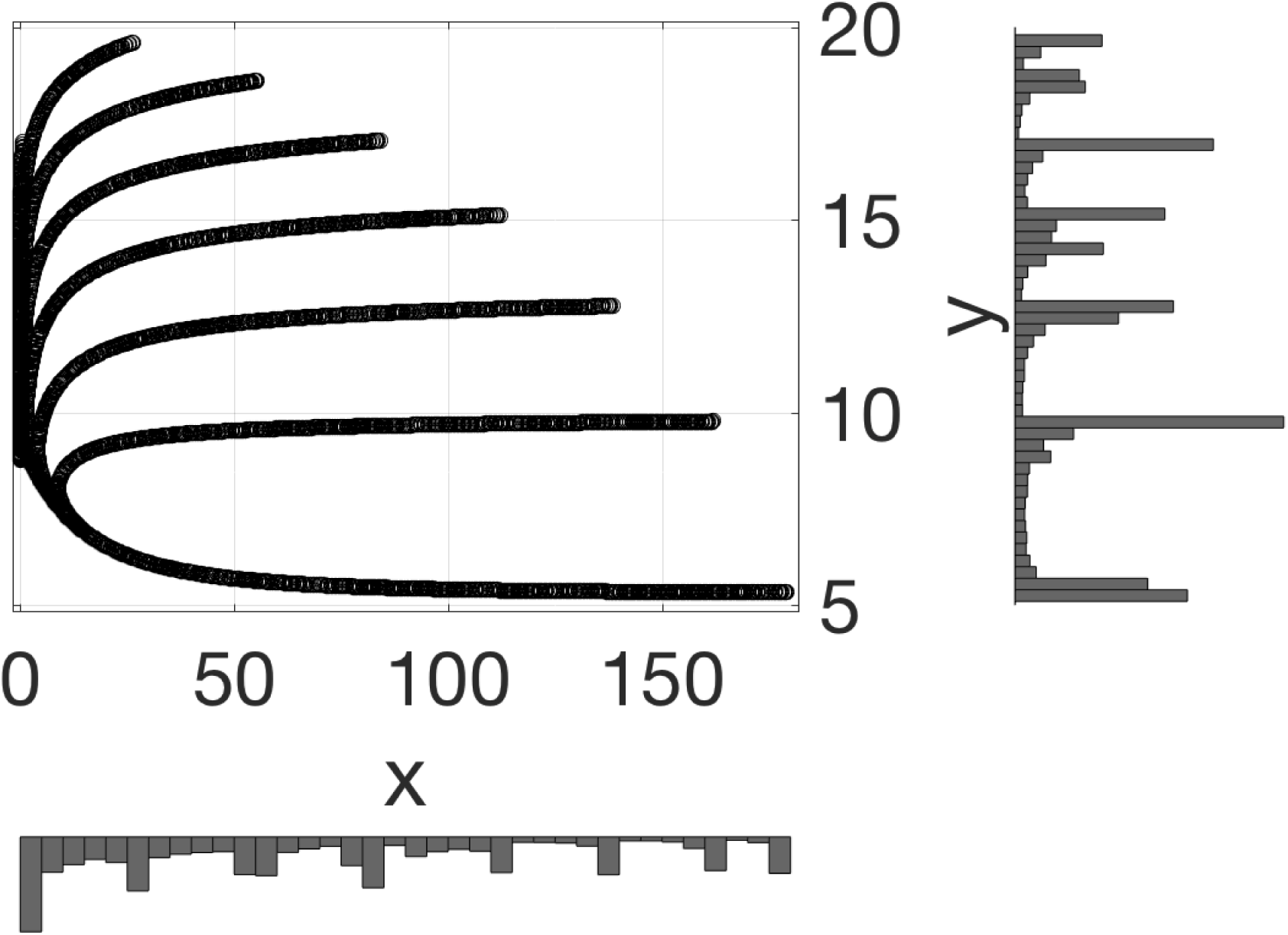
Scatter plot and marginal distributions for the unity model (1) wight *T* = 40.

**Figure SI-3.3:**
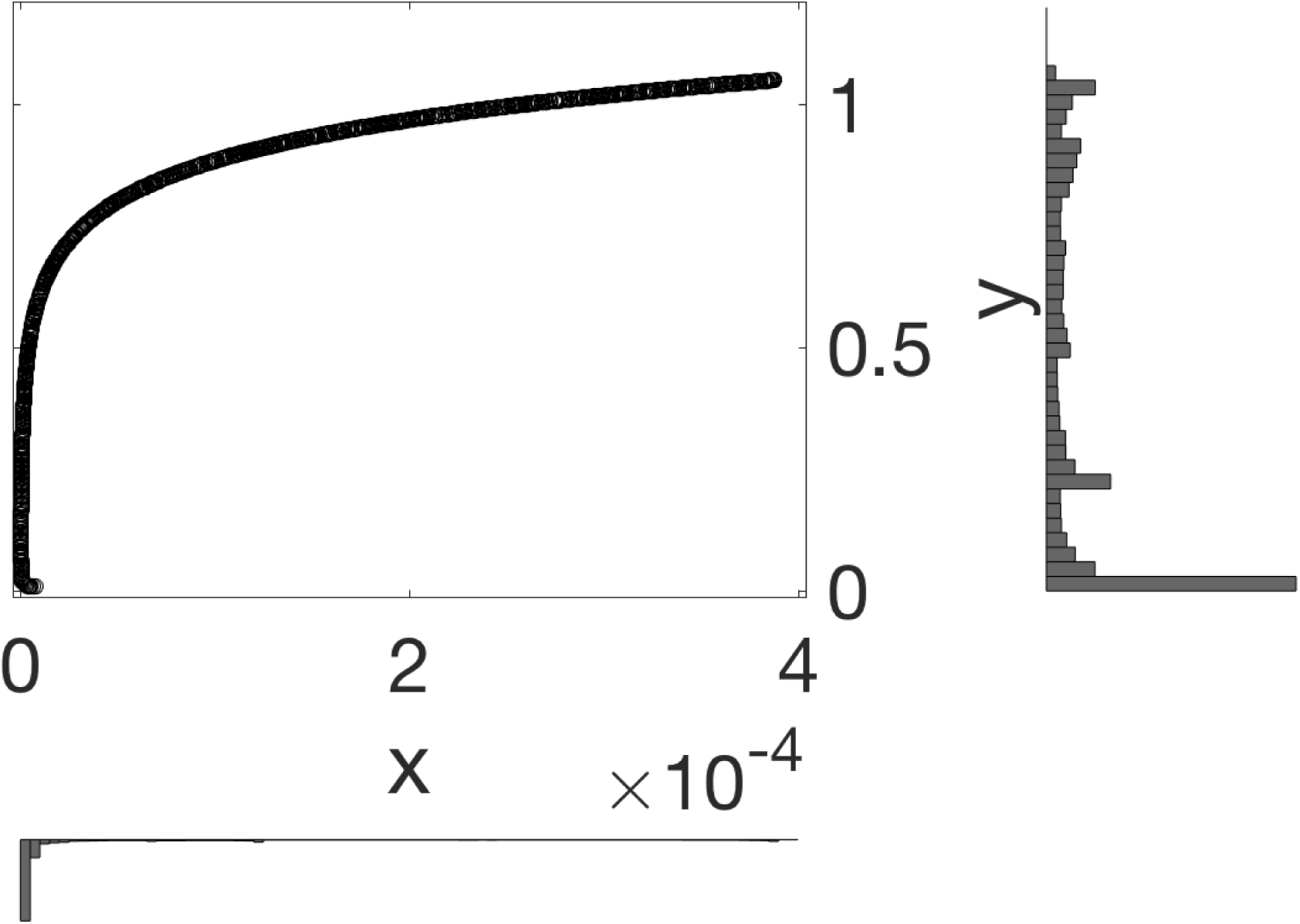
Scatter plot and marginal distributions for the FCD model (2) whit *T* = 5.

### SI-4 Bifurcation trees with respect to the parameter *m*

Fig. SI-4.1 demonstrates examples of bifurcation diagrams, when the values of the parameter *m* are allowed to vary: (*a*) the leftmost column plots correspond to a strange attractor in the model (1) with fixed parameter values *σ*_*x*_ = *σ*_*y*_ = 10^4^, *θ*_*x*_ = 10, *θ*_*y*_ = 1, *µ*_*x*_ = 100, *µ*_*y*_ = 10, *M*_*x*_ = *M*_*y*_ = 1, *T* = 1, and ∆ = 10^−4^; (*b*) the middle column plots correspond to the unity model (1) with *T* = 100 and ∆ = 10^−2^ (and all other unit parameter values also kept fixed); and (*c*) the rightmost column plots correspond to a strange attractor in the FCD model (2) with *a* = *Y*_0_ = *c* = *d* = 1.0, *T* = 5, and ∆ = 0.2.

**Figure SI-4.1:**
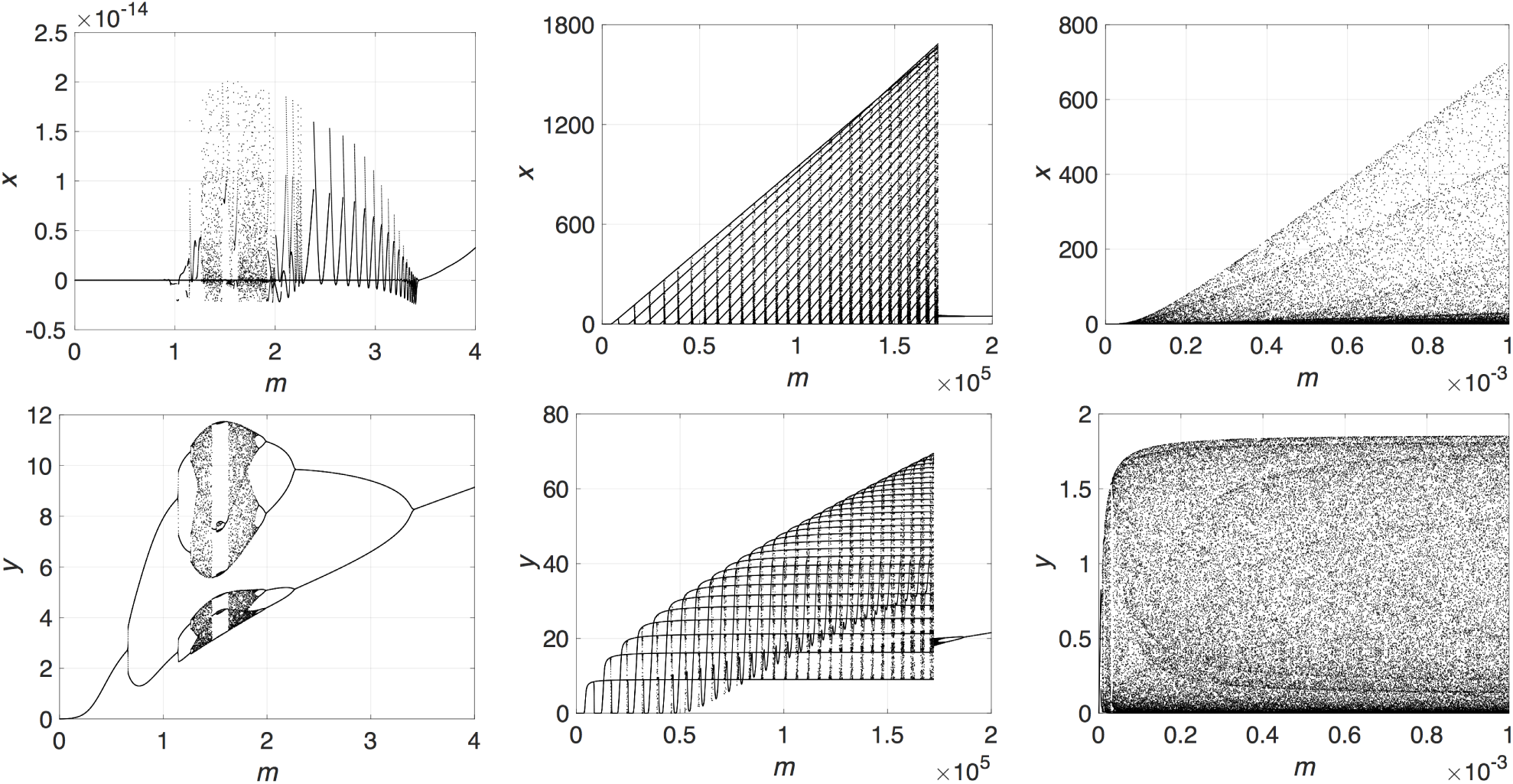
Bifurcation diagrams. The panels in the left column correspond to the model (1); the panels in the middle column correspond to the unity model obtained from the model (1) by setting unit values to all model’s parameters; and the panels in the right column correspond to the FCD model (2).

### SI-5 Experimental results

In [14], experiments were performed on intact *C. elegans* worms in microfluidic chambers, measuring the response of odor-sensing AWA neurons (quantified by intracellular Ca^2+^ activity as measured by an AWA-specific GCaMP sensor) to periodic on-off pulses of diacetyl. Shown in Fig. SI-5.1 is a harmonic response in one experiment to a pulse with period *T* = 39s, as well as, for another experiment, what look like sub-harmomic responses when the period of pulses is shorter, *T* = 15s. The experiments used technology developed in [54]. (Not shown are preparatory odor pulses, used to calibrate the recordings across experiments by waiting for stabilized responses.)

### SI-6 Verification of smoothness of Φ(*z, α*)

We wish to prove that Φ(*z, α*) is smooth, for our models with the piecewise defined input (3). The result follows from a general theorem on the smooth dependence of the solutions of the given ODE on initial conditions and parameters, see e.g. [55]. Indeed, let us first represent Φ(*z, α*) in the following superposition form,

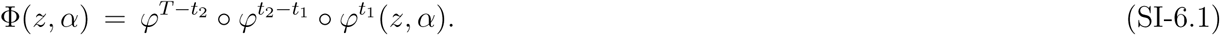

Let *z*_1_(*z, α*) = *φ*^*t*_1_^(*z, α*), *z*_2_(*z*_1_,*α*) = *φ*^*t*_2_−*t*_1_^(*z*_1_,*α*), and *z*_3_(*z*_2_,*α*) = *φ*^*T*−*t*_2_^(*z*_2_, *α*).

Since *z*_1_(*z, α*) smoothly depends on *z, z*_2_(*z*_1_) smoothly depends on *z*_1_ (viewed as an independent initial condition), and, analogously, *z*_3_(*z*_2_, *α*) smoothly depends on *z*_2_, it follows from the differentiation chain rule applied to the superposition (SI-6.1) that Φ(*z, α*) smoothly depends on the state variable *z* [55].

Similarly, because each of the shift maps, *φ*^*T*−*t*_2_^(*z*_2_, *α*), *φ*^*t*_2_−*t*_1_^(*z*_1_,*α*), and *φ*^*t*_1_^(*z, α*), smoothly depends on the parameter *α*, we can conclude that Φ(*z, α*) as well smoothly depends on the parameter *α* [55].

Moreover, it can be proved that Φ(*z, α*) is a diffeomorphism [55].

**Figure SI-5.1:**
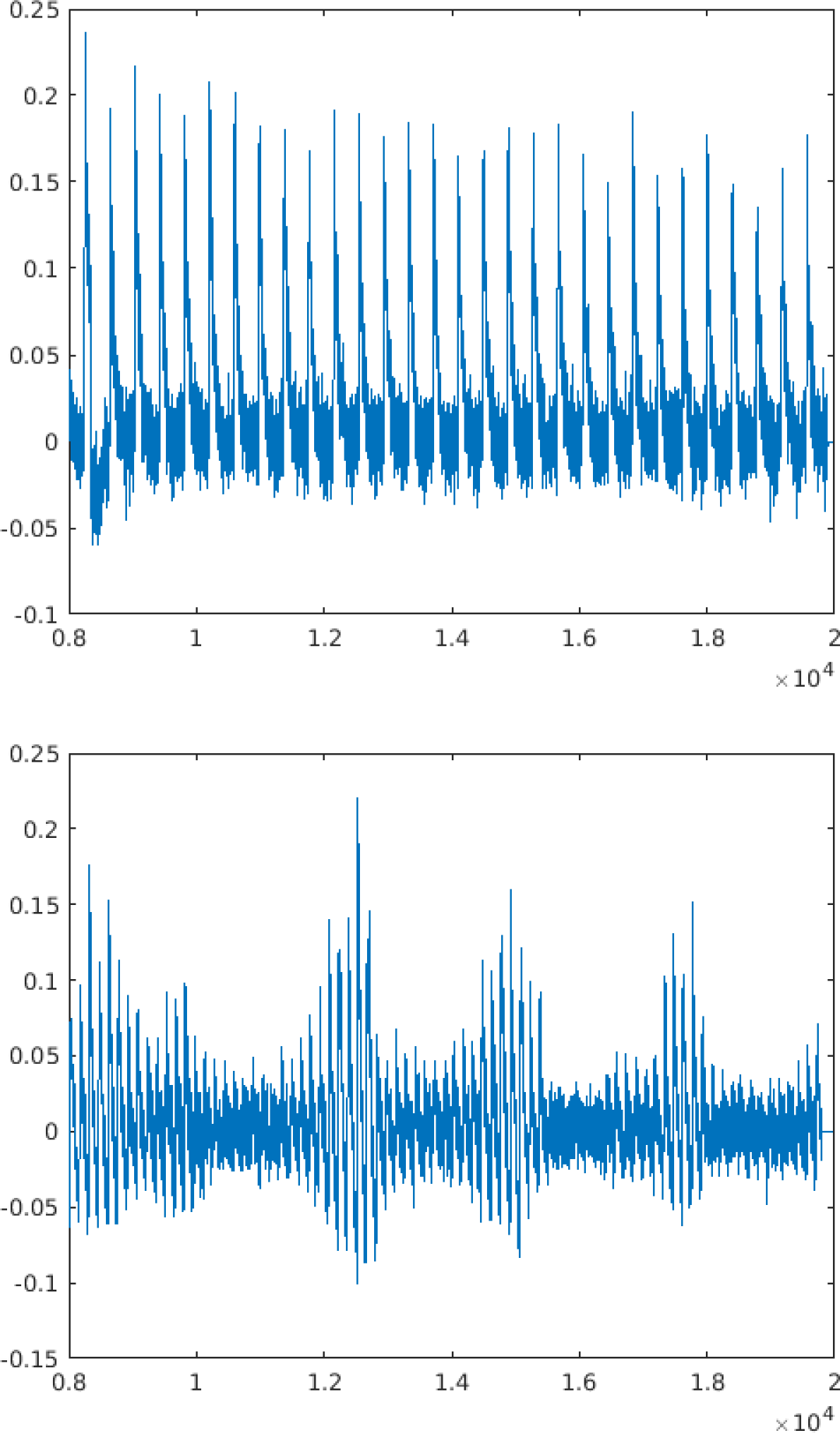
Transition from entrainment to subharmonics, as period decreases, in a worm odor-sensing experiment. Two selected responses to longer and shorter periods, from the work [14]. The *x*-axis represents time in units of 0.1 seconds, and the *y*-axis is intracellular Ca^2+^ activity in arbitrary units. Top: A trace showing an approximately harmonic (entrained) response to a pulse train with period *T* = 39s. Bottom: A trace showing a non-entrained response, with an apparent lower-frequency component, to a pulse train with period *T* = 15s. Pulse duraction ∆ = 10s in both cases. See [14] for details.

